# Decreased inter-hemispheric connectivity predicts a coherent retrieval of auditory symbolic material in a laboratory model of cultural transmission

**DOI:** 10.1101/2023.06.07.543882

**Authors:** Leonardo Bonetti, Anna Kildall Vænggård, Claudia Iorio, Peter Vuust, Massimo Lumaca

**Affiliations:** Center for Music in the Brain, Department of Clinical Medicine, Aarhus University & The Royal Academy of Music, Aarhus, Aalborg, Denmark; Centre for Eudaimonia and Human Flourishing, Linacre College, University of Oxford, Oxford, United Kingdom; Department of Psychiatry, University of Oxford, Oxford, United Kingdom; LEAD-CNRS UMR 5022, Université de Bourgogne, 21000 Dijon, France

**Keywords:** Asymmetry, information flow, brain connectomes, hemispheric specialization, signalling games, cultural transmission

## Abstract

Investigating the transmission of information between individuals is essential to understand how human culture evolved. Coherent information transmission (i.e., transmission without significant modifications or loss of fidelity) helps preserving cultural traits and traditions over time, while innovation may lead to new cultural variants. Although much research has focused on the cognitive mechanisms underlying cultural transmission, little is known on the brain underpinnings of coherent transmission of information. To address this gap, we combined a laboratory model of cultural transmission, the signalling games, with structural (from high-resolution diffusion imaging) and functional connectivity (from resting-state functional magnetic resonance imaging [fMRI]). We found that individuals who exhibited more coherence in the transmission of the information were characterized by lower levels of both structural and functional inter-hemispheric connectivity. Specifically, higher coherence negatively correlated with the strength of bilateral structural connections between frontal and subcortical, insular and temporal brain regions. Similarly, we observed increased inter-hemispheric functional connectivity between inferior frontal brain regions derived from structural connectivity analysis in individuals who exhibited lower transmission coherence. Our results suggest that inter-hemispheric connections may bwe detrimental for preserving coherence in information transmission, while a certain degree of lateralization in the brain may be required.

## Introduction

Social learning, information transmission, and innovation are key elements in human cultural evolution and play a significant role in the development and variation of symbolic systems like language and music. For example, musical information is typically passed down from one generation to the next through oral or written transmission (Patterson, 2015). This process can happen without significant modification or loss of fidelity, maintaining the music very similar to that of the previous generations. Conversely, this process can also lead to differences which add novelty to the music of the next generation. In the former case, the transmission of information is considered coherent, and the musical tradition is preserved, while, in the latter, coherence is lost, and novel musical features are introduced. Accordingly, Tomasello (1999) highlighted that accurate social learning and coherence during transmission of information help preserving cultural traits and traditions. On the contrary, imitation errors in the transmission of information which are maintained in the following generation lead to new cultural variants (Bentley & O’brien, 2011; Boyd & Richerson, 1988; Caldwell et al., 2016). For this reason, the innovation introduced by imitation errors in the context of cultural transmission explains some of the diversity observed within the same culture over time or between different cultures (Fitch, 2011; Rzeszutek et al., 2012).

Although the transmission of information and symbolic messages across individuals of a culture over time is a complex phenomenon, simplified yet accurate models of cultural transmission can be achieved in controlled experimental settings. One such model is the signalling game in which a sender and a receiver must coordinate through several interactions to a common ‘code’ (Lewis, 2008; Skyrms, 2010). In this setting, a code is a system of signal-meaning (state) mappings that exists in the mind of the communicators (sender and receiver) wherein a certain signal matches a certain meaning. In each signalling trial, the sender has private access to a meaningful state and uses a signal to inform the receiver on the identity of the state. The receiver must in turn act and, if the action matches the observed state, communication is successful. During the game, the sender can partly adapt their mappings to the ones used by the receiver and vice versa. Hence, information flow can be bi-directional or shifted toward the sender or the receiver. Previous studies showed that when senders and receivers have fixed roles (Moreno & Baggio, 2015), signalling games turn from a model of coordination to a model of cultural transmission (Lumaca & Baggio, 2017, 2018; Nowak & Baggio, 2016). Critically, this model can serve useful to assess the coherence of the transmission of the information during the communication between the sender and the receiver. This measure is known as transmission coherence and, in simple terms, means that if the sender is coherent in sending the signals and their mapping to meanings throughout trials, transmission coherence will tend to be high.

Previous research on the neural underpinnings of transmission biases has been conducted in a series of signalling game studies using electroencephalography (EEG) (Lumaca & Baggio, 2016; Lumaca, Haumann, et al., 2018) (For a review)(Lumaca, Ravignani, et al., 2018). First, authors linked cultural transmission of melodic signals to brain responses indexing sensory memory and detection of environmental irregularities, discovering that their MMN latency predicted the learning, transmission, and structural modification of the signalling systems learnt by the participants (Lumaca & Baggio, 2016). Lumaca, Haumann, et al. (2018) replicated this finding to the rhythmic dimension of music. They discovered that rhythmic transmission and regularization of short tone sequences could be predicted by the MMN latencies measured in a temporal oddball paradigm. The authors suggested that isochronicity in communication systems originated in neural constraints on information processing, a phenomenon which may then be expressed and amplified during cultural transmission. The relationship between neural data and cultural transmission have been further explored by using tools such as functional magnetic resonance imaging (fMRI) and brain connectivity (M. Lumaca et al., 2021; Lumaca, Kleber, et al., 2019). In Lumaca, Kleber, et al. (2019), the authors used resting-state functional connectivity (rs-FC) to assess the relationship between features of the functional brain networks during rest and learning and transmission of an artificial tone system. The rs-FC measure is derived by the statistical correlation of the spontaneous slow neural fluctuations of their associated hemodynamic responses (<0.1 Hz) observed between separate brain areas and estimates their degree of communication (Bruzzone et al., 2022; Deco & Kringelbach, 2014; Deco et al., 2015; Lord et al., 2017; Massimo Lumaca et al., 2021; Lumaca, Kleber, et al., 2019; Lumaca et al., 2022). The authors found that the inter-hemispheric functional connectivity within fronto-temporal auditory networks during resting state predicted learning, transmission, and structural modification of the artificial tone system used in the game. Afterward, M. Lumaca et al. (2021) investigated measures of brain structural connectivity and linked them to signalling games. As opposed to its functional counterpart, structural connectivity is often derived from diffusion tensor imaging (DTI) and measures the strength of the physical (axonal) connections that link the brain regions and form the biological infrastructure for neuronal signalling and communication (Van Hecke et al., 2016). Stronger connections and higher fiber density can be measured by different parameters such as fractional anisotropy (FA) and used as proxies of the quantity of information that is potentially carried from one region to another. In the context of cultural transmission, M. Lumaca et al. (2021) collected DTI data and discovered that FA of auditory callosal pathways predicted the fidelity of transmission of the artificial tone system used in their experiment. The results of the previous two studies suggest that cultural transmission relies on the distribution of neural sources between left and right auditory cortex. This hypothesis was explored more deeply in a new study looking at the relationship between transmission behaviours and left-right neuroanatomical auditory asymmetries (Lumaca et al., 2023). They showed that high accuracy in the transmission of an artificial tone system across generations was linked to reduced rightward asymmetry of cortical thickness in the Heschl’s sulcus.

These studies advanced our understanding of the neural underpinnings of cultural transmission. However, they were mainly limited to the investigation of the auditory cortex, while a broader network is most likely involved in a complex and stratified process like cultural transmission of symbolic codes (Lumaca et al., 2022). Along this line, previous studies have investigated the neural correlates of a plethora of cognitive processes which are arguably involved in cultural transmission, such as auditory processing, encoding and recognition of information, creativity, and decision-making. For instance, several studies showed the key role of Heschl’s and superior temporal gyri and insula in auditory processing (Brattico & Pearce, 2013; Mutschler et al., 2007). Other works highlighted the involvement of medial temporal lobe and hippocampus for several memory systems (Bird, 2017; Bonetti, Brattico, et al., 2022; Bonetti et al., 2020; Leonardo Bonetti et al., 2021; L Bonetti et al., 2021; Bonetti, Carlomagno, et al., 2022; Brown & Aggleton, 2001; Fernandez-Rubio et al., 2022; Fernández-Rubio et al., 2022). Similarly, decision-making and creativity have been associated with a complex interplay occurring between multiple brain regions including the prefrontal cortex, anterior cingulate, basal ganglia, hippocampus, and amygdala. (Clark & Manes, 2004; Lockwood & Wittmann, 2018; Neubert et al., 2015).

Taken together, these studies provided evidence of associations between cultural transmission and measures of brain structure and functioning. However, much remains to be discovered. For instance, to our knowledge, no evidence has ever been reported on the relationship between neural data and coherent transmission of information. However, this is highly important for two complementary reasons: 1) to understand the neural substrate that has allowed individuals to preserve information and cultures, and 2) to investigate how cultural diversity evolved. Thus, in this study, we aimed to assess the relationship between the degree of coherence in information transmission in signalling games and the inherent organization of connectivity patterns of the participants.

## Methods

### Subjects

Fifty-two right-handed volunteers (33 females, mean age 24.5 years, range 20-34) were recruited for the experiment. Participants with neurological or psychiatric disorders, as well as hearing impairments, were excluded. One participant (female) participated in only one session of the experiment; thus, the experiment was completed by 51 participants. All participants were non-musicians, with no more than six years of formal musical training (mean = 0.6 years, standard deviation = 0.98) (Zhang et al., 2020). All subjects provided written consent and completed an MRI safety form before entering the MR scanner. The Central Denmark Region’s ethical committee (nr. 1083) approved the study procedure.

### Study Design

In this study, the participants completed two experimental sessions (one MRI and one behavioural) on two separate days (mean 20 days apart, range 13-30 days). The MRI session included four distinct scans: resting-state fMRI, high-resolution structural (MP2RAGE), fMRI (T2*-weighted), and HARDI (see Session 1: MR acquisition). Following debriefing, participants completed a digit span test (Orsini et al., 1987) (backward and forward) to assess their working memory and attention capacities. Participants also completed the Goldsmiths Musical Sophistication Index (Gold-MSI) questionnaire, which measured their musical background. Digit span and Gold-MSI scores are not associated to behavioural performance in signalling games (Lumaca, Kleber, et al., 2019).

In the second session, participants took part in two consecutive signalling games (**Figures 1 and 2**) (Moreno and Baggio 2015; Nowak and Baggio 2016) (see Session 2: Signalling Games). In the first game (Game 1), the participant played as receiver (learner) of an artificial tone system (the seeding material or “seed”) that consisted of 5-tone auditory patterns, each conveying a particular emotion (a facial expression). Participants were instructed to learn the seed from an experimenter’s associate (confederate). The same participant played as the sender (transmitter) in the second game (Game 2) and was asked to reproduce the seed learnt in Game 1 by memory. The two games were played by different experimenters’ associates (N=3). This study only reports on the analysis of dMRI and resting-state functional imaging data in relation to the performance of participants in Game 2. The analysis of the fMRI oddball experiment is outside the scope of this paper and is detailed elsewhere (Lumaca, Kleber, et al., 2019).

**Figure 1.**
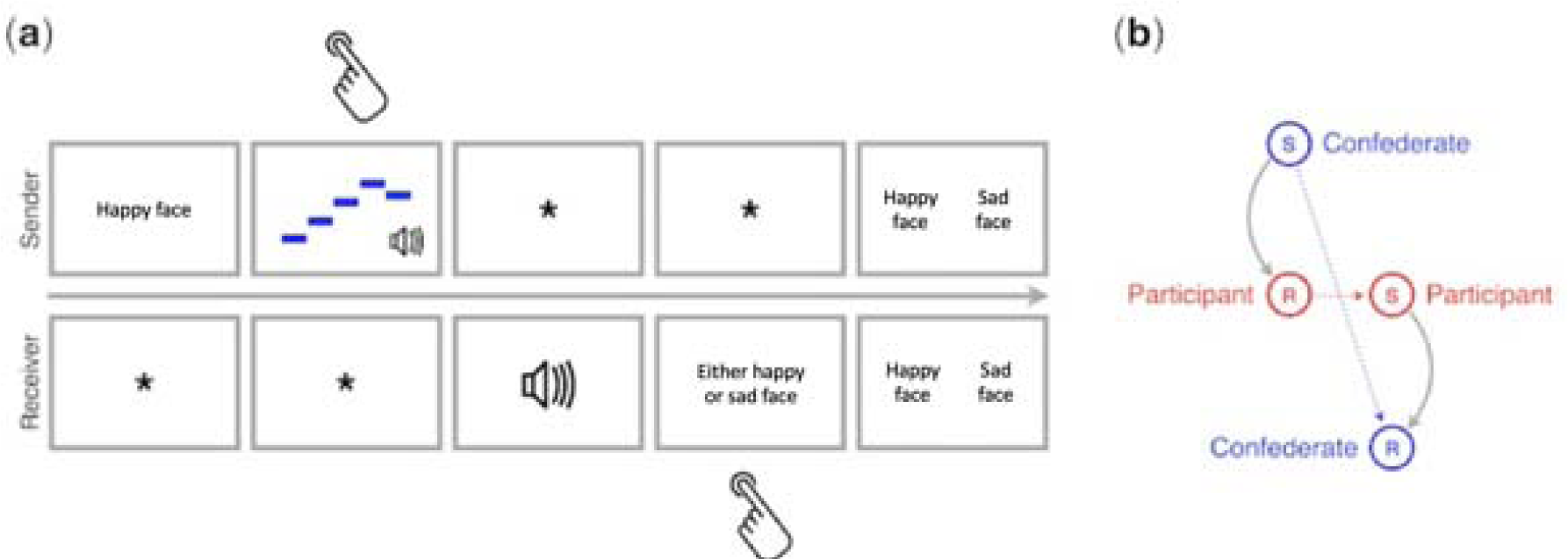
Example of a trial from the signalling games (left panel) played by participants on the second session of the study and experimental transmission design (right panel). **(a)** The top and bottom rows show what the sender and receiver saw on their screens, respectively. The task for the sender was to compose a five-tone sequence to be used as a signal of the simple or compound emotion expressed by the face presented on the screen at the start of the trial, and for the receiver to respond to that signal by choosing the face that the sender had seen. The sender and the receiver converged over trials on a shared mapping of signals (tone sequences) to meanings (emotions). Hand symbols indicate when the sender and the receiver had to produce a response. Feedback was given to both players simultaneously, displaying the face seen by the sender and the face selected by the receiver in a green frame (matching faces; correct) or in a red frame (mismatching faces; incorrect). Time flows from left to right. **(b)** A diagram of the signalling games played by the participant is also shown. The participant played as receiver (R) with a confederate of the experimenters playing as sender (S) in Game 1. Roles switched in Game 2.

**Figure 2.**
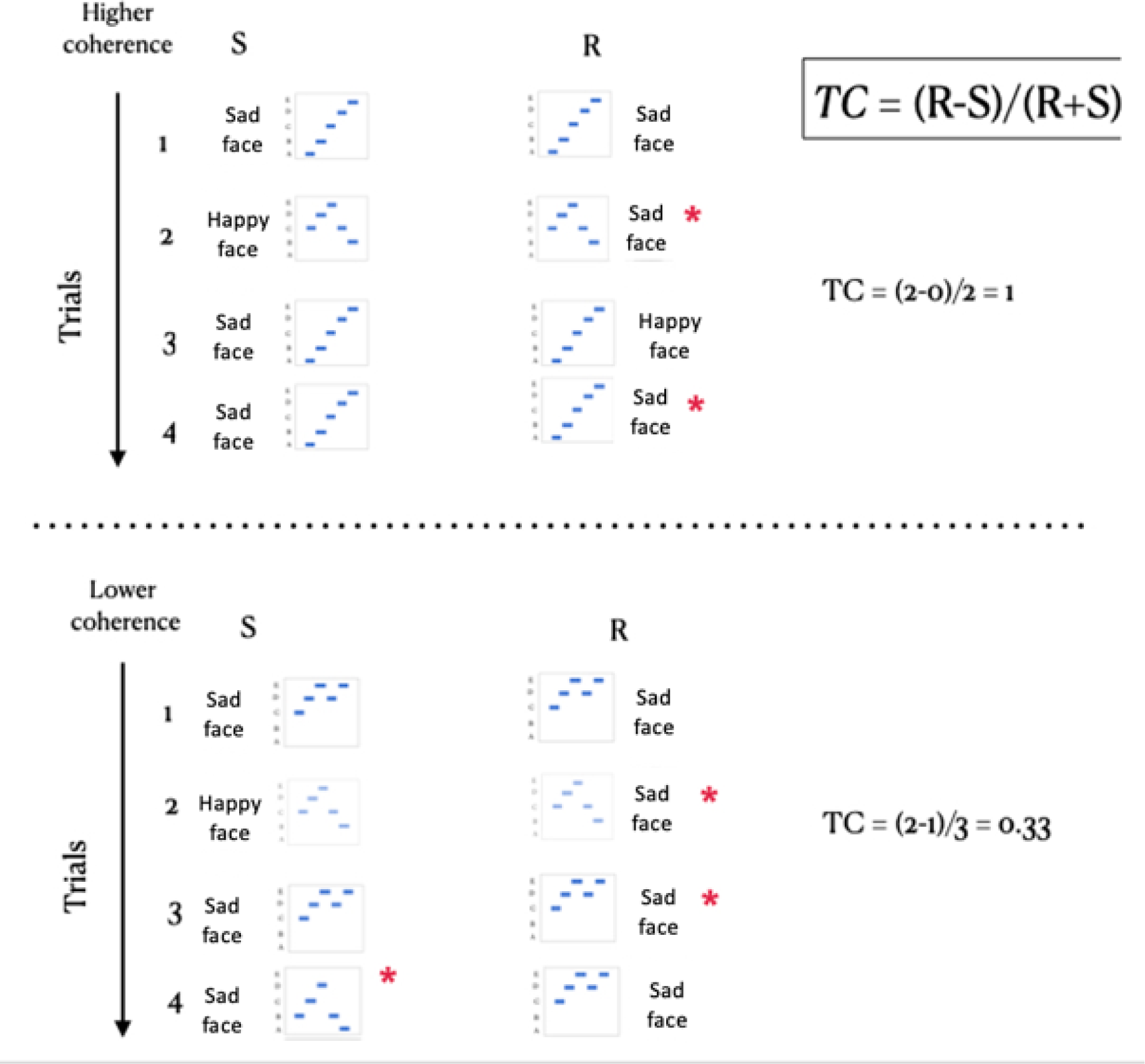
An example on how transmission coherence was calculated in high consistent (top panel) and low consistent trials (bottom panel). Transmission coherence (TC) is defined as the number of code changes made by the receiver minus the number of code changes made by the sender, over the total of code changes. It ranges from 1 (all code changes are made by the receiver) to -1 (all code changes are made by the sender). Positive TC indicates that the sender in game 2 has been highly consistent in sending the same signal for a given emotion throughout the signalling trials (top panel). Lower TC values indicate that the sender was less consistent in performing that task (bottom panel). The asterisk indicates a mapping change.

### Signalling games

Signalling games (Lewis, 2008; Skyrms, 2010) are a type of games that model social coordination in language and other human symbolic systems, either in laboratory settings or in computational simulations. These games typically consist of two players, a sender and a receiver, who engage in several rounds of signalling interactions. The goal of the players is to establish a common code, an artificial language system made up of signals that represent events. In each round of the game, the sender observes a state from a set of possible states and selects a signal to send to the receiver that represents that state. The receiver then chooses an action that should correspond to the initial state. Both players receive (the same) feedback at the end of each round. Positive feedback is given if the identity of the state observed by the sender and chosen by the receiver match (correct trial), while negative feedback is given otherwise. Information flow can be captured in chains of two or more players (Skyrms, 2009), where the sender may consistently use a signal for one state throughout multiple trials. In turn, the receiver may adapt the signal-state mappings to coordinate on a common code, and the net information flow is from sender to receiver. Moreno and Baggio (2015) demonstrated that this condition is achieved when the sender and receiver play with fixed roles throughout the game. A net information flow from sender to receiver is a key property of cultural transmission and cumulative cultural evolution.

### Stimuli

The stimuli used in this study consisted of 5-tone auditory sequences generated by the sender using the computer keyboard (see “Transmission Design” below). Each pure tone had a duration of 50 milliseconds and was drawn from the equal-tempered version of the Bohlen-Pierce (BP) scale (Mathews et al., 1988). The BP scale is a macrotonal tuning system in which a just twelfth (“tritave”) is divided into 13 logarithmic even steps. Participants were unable to use prior musical knowledge to map signals to meanings (“focal bias”), as they were unfamiliar with this scale. The frequency of each tone was determined by the formula F = k * 3 (n/13), where n represents the number of steps along the scale, and k is a constant corresponding to the fundamental frequency. In this study, k was set to 440 Hz, and n was set to 0, 4, 6, 7, and 10 (to maximize the low-integer frequency ratio of tone combinations). The signals were delivered via headphones at a loudness level of 70 dB. The states, which were denoted by the signals, consisted of five emotion categories of varying complexity, including simple emotions (peace, joy, sadness) and compound emotions (peace x joy, peace x sadness). The states were presented on the screen as facial expressions of one actor.

### Transmission design

This study employed a transmission design that consisted of two consecutive fixed role signalling games, referred to as iterative signalling games (Lumaca & Baggio, 2016). During Game 1, the participant acted as the receiver and was instructed to learn an artificial tone system, or “seed”, that consisted of 5-tone sequences (signals) mapped to facial expressions (states), sent by the confederate (sender). The seed was originally devised by the experimenter and controlled of its melodic contour complexity. In the Game 2, participants switched their player role to sender and were asked to transmit the code learned to a new confederate (receiver), regardless of the outcome. Each participant was assigned to one of three ‘seed’ groups (**Figure 1**). Each seed consisted of two high-entropy signals (Shannon Entropy, H = 1 bit), two low-entropy signals (H = 0.81 bits), and a monotone signal (H = 0 bits). Pseudo-randomization of signal-meaning mappings was utilized in the seed to prevent real-world associations that could be exploited by participants to reproduce the code (e.g., signals with upward melodic lines denoting positive-valence states).

At the beginning of a trial, the sender observed a facial expression for a duration of three seconds and was then asked to produce a five-tone sequence signal denoting that state. The signal was produced using the number digits (1-5) of the computer keyboard, with each key mapped to one BP tone. The sequence could be tried with no limits of time before being sent to the receiver. Upon hearing the signal via headphones, the receiver used the number pad (1-5) to select the facial expression that matched the initial state, with duration being self-paced. Feedback was presented to both players for a duration of three seconds, indicating whether the expression chosen by the receiver matched the one observed by the sender. The game ended after a fixed number of trials were completed (70 trials in Game 1 and 30 trials in Game 2).

### Transmission coherence

Transmission coherence (TC) was measured during Game 2. It quantifies the consistency with which each participant (sender) produces the same signal/mapping for a given state throughout the 30 signalling trials, regardless of the original code (i.e., the seed). As shown in **Figure 2**, it is defined as the number of changes (in mappings) made by the receiver (R) minus the number of changes made by the sender (S), over the total of code changes: TC = (R-S)/(R+S). It ranges from 1 (the sender is fully consistent across trials with their code production) to -1 (the sender is fully inconsistent and changes the mapping from trial to trial).

### MR acquisition

Brain scans were collected using a 3T MRI scanner (Siemens Prisma). During the fMRI resting-state scan, a total of 600 volumes was acquired using fast T2*-weighted echo-planar imaging (EPI) multiband sequence (TR, 1000 ms; TE, 29.6 ms; voxel size, 2.5 mm3). In the meanwhile, participants were asked to maintain their eyes closed and avoid mind wandering. We used an MP2RAGE sequence for the acquisition of a high-resolution T1-weighted (T1w) image (TR=5000 ms; TE=2.87 ms; voxel size=0.9 mm3) (10 min). Finally, we collected high-angular-resolution diffusion (HARDI) images with the following acquisition parameters: 90 diffusion directions at b=2800 s/mm2; 60 directions at b=1200 s/mm2; 30 directions at b=700 s/mm2; 11 directions at b=5 s/mm2, with the different b-shells acquired in the same series (flip angle=90°, TR/TE=2972/65 ms, voxel size=1.8 ⨉ 1.8 ⨉ 1.8 mm, matrix size=112 x 112, number of slices=90). The phase-encoding direction was anterior to posterior (AP). An opposite phase-encoding direction (i.e., PA) was also acquired (b=5 s/mm) to allow EPI distortion correction (Andersson et al., 2003; Holland et al., 2010).

### Structural connectome

The MRtrix3 software package (version 3.0_RC3) was employed to pre-process and analyse HARDI images (Tournier et al., 2019)(http://mrtrix.org). Additionally, other software packages were utilised in this study, including FSL (Jenkinson et al., 2012) for DWI denoising and 4-tissue segmentation of T1w images, ANTS N4 (Avants et al., 2014) for DWI bias field correction, SPM12 (r7487) implemented in Matlab R2016b (Mathworks) for background noise-cleaning of T1w MP2RAGE, ROBEX for brain extraction of T1w images (Iglesias et al., 2011), and FreesSurfer (freeSurfer-Linux-centos6_x86_64-stable-pub-v5.3.0) for the pre-processing and grey-matter parcellation of T1w images (Fischl, 2012). **Figure 3** (top panel) illustrates a schematic of the dMRI connectomic pipeline. The processing for each subject involved:

- Pre-processing of DWI images. This step involved their denoising (Veraart et al., 2016), unringing (Kellner et al., 2016), motion and distortion correction (Andersson & Sotiropoulos, 2016), and B1 bias field inhomogeneity correction (Tustison et al., 2010). From pre-processed dMRI images we obtained 3-tissue response functions (RFs) that represented single-fiber white matter (WM), grey matter (GM), and cerebrospinal fluid (CSF), using an unsupervised approach (Dhollander et al., 2016). Fiber orientation distributions (FODs) were then computed at the single-voxel level using multi-tissue Constrained Spherical Deconvolution (CSD) on average response functions (Jeurissen et al., 2014). Lastly, global intensity normalization of the 3-tissue parameters was performed (Raffelt et al., 2017). The computation of group-average RFs and global intensity normalization are necessary steps for inter-subject connection density normalization.
- Pre-processing and cortical parcellation of T1w images. T1w MP2RAGE images were cleaned from background noise (Marques et al., 2010), followed by brain extraction (Iglesias et al., 2011; O’Brien et al., 2014). T1-to-DWI intermodal registration was performed (Bhushan et al., 2015), and segmentation of T1w into cortical grey matter, subcortical grey matter, white matter, and CSF was conducted (Patenaude et al., 2011; Zhang et al., 2001). This step is necessary for anatomically constrained tractography (ACT). The FreeSurfer processing pipeline (default) (Dale et al., 1999) was employed for cortical parcellations of T1w images, which resulted in 148 cortical regions (74 homologs) based on the Destrieux atlas (Destrieux et al., 2010), plus 17 regions based on FreeSurfer segmentation (7 subcortical homologs, bilateral cerebellum, and midbrain).
- Tractography. The whole-brain fiber tractography was conducted using the iFOD2 probabilistic algorithm with dynamic seeding (20 million seeds) in MRtrix3. The ACT algorithm with the ‘backtracking’ option was employed to apply biological tissue priors to streamline generation, ensuring that streamlines only terminated at anatomically correct tissue locations (Smith et al., 2012). To determine a weight for each streamline, the SIFT2 algorithm (Smith et al., 2015) was used on the tractogram. The SIFT2 weights provide the tractogram with quantitative and more biologically meaningful properties.
- Construction of the structural connectomes. The tractogram streamlines were assigned to nodes (anatomical regions), which were defined by the Destrieux parcellation, resulting in an undirected N ⨉ N matrix (where N is the number of nodes) (Smith et al., 2015). Based on fiber “connection density” (or fiber bundle capacity, FBC), one connectome (165 ⨉ 165) was constructed, where each entry represented the strength of connection between any two areas of the parcellation. To calculate FBC for each edge, the sum of SIFT2 weights was multiplied by a scalar proportionality coefficient ‘μ’ (Smith et al., 2020).

**Figure 3.**
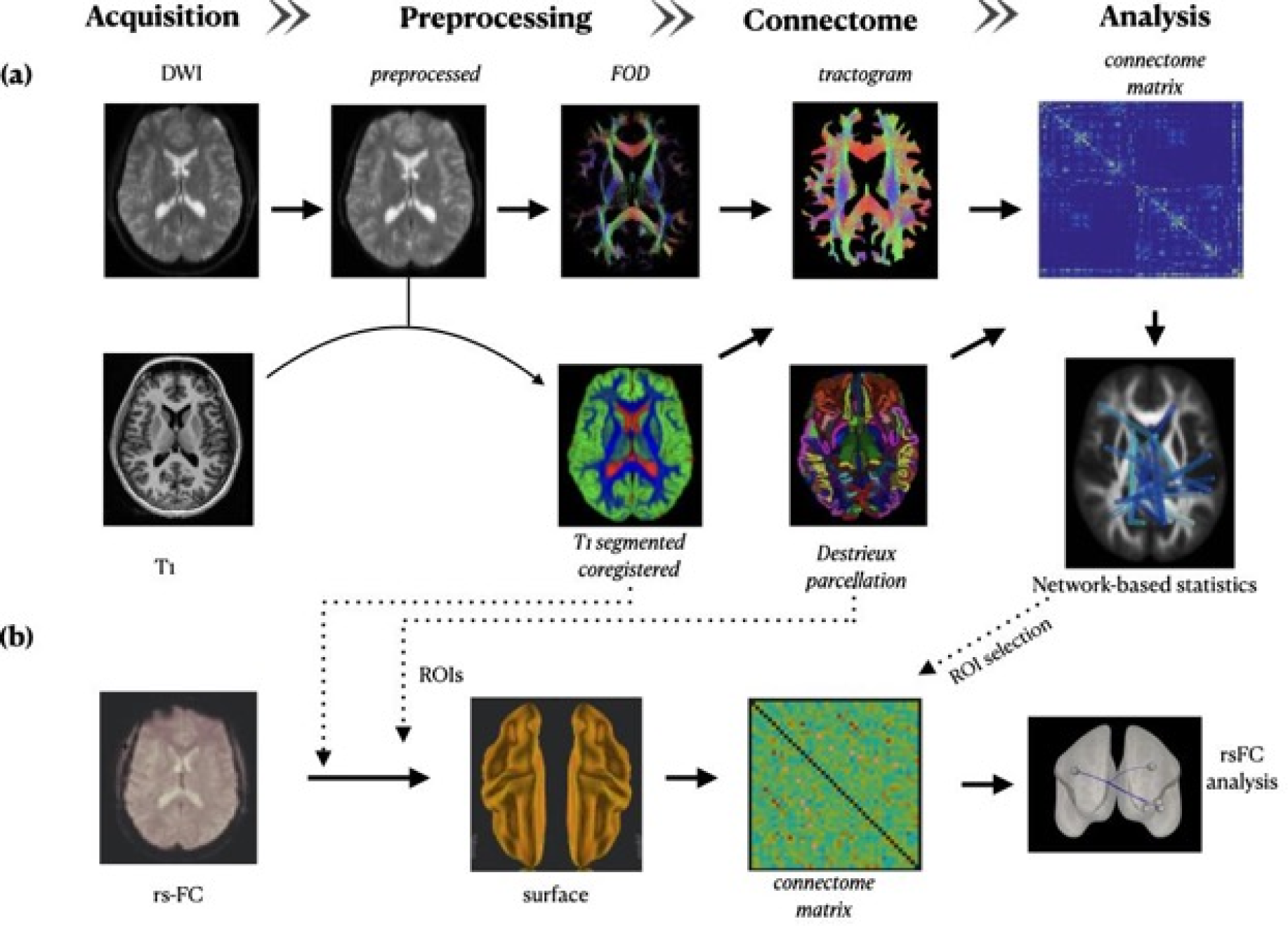
The figure depicts a schematic of the MRI connectomic pipeline used for surface-based structural and functional connectome analyses. **(a)** For each individual, we processed high angular resolution diffusion (HARDI) images to generate fiber orientation distributions (FODs) (see Methods for more details). High-resolution T1 images (MP2RAGE) were co-registered to pre-processed DWI images and then segmented into four tissues (cortical grey matter, subcortical grey matter, white matter, and CSF). The combination of T1 segmented images and FODs images allowed the use of anatomically constrained tractography (ACT) for the construction of whole-brain SIFT2-based tractograms (20 million streamlines). These tractograms were combined with the Destrieux parcellation (165 x 165) of MP2RAGE images generated with Freesurfer, to obtain the anatomical connectome matrix. This matrix contains fiber bundle capacity (FBC), a measure of connectivity strength that reflects how much information each connection can carry from one brain region to another. Non-parametric threshold-free network-based enhanced statistics (TFNBS) (number of permutations = 5000) was used on interregional connectivity matrices to find an anatomical network (N=48 nodes) where FBC values were correlated with TC scores. Finally, from this anatomical network, we extracted a core subnetwork of the 20% highest-degree regions-of-interest (ROIs) that was used to perform rs-FC analyses. **(b)** rs-FC images were pre-processed and co-registered to T1W segmented images, then converted to surface level space and parcellated based on the Destrieux atlas. Blood oxygenation level dependent (BOLD) signals were extracted from areas of the core network (N=9 nodes). Pearson’s R-values were calculated, Z transformed, and analysed using non-parametric cluster-based analyses (spatial pairwise clustering, SPC; number of permutations = 1000) to examine in which cluster of connections Fisher’s z-transformed was associated with TC scores.

### Functional connectome

The SPM CONN toolbox (Whitfield-Gabrieli & Nieto-Castanon, 2012) was utilized for processing and analyses of resting-state functional connectivity (rs-FC) (see Fig. 3 bottom panel). The processing for each subject involved:

- Pre-processing of rs-fMRI images. The rs-FC images underwent several pre-processing steps, including realignment and unwarping, ART-based outlier detection (motion correction = 0.9 mm; Global-signal z-value threshold = 5), direct co-registration to T1w segmented images, conversion from volume-level to surface-level data by resampling functional data to fsaverage space, iterative diffusion smoothing with a series of 40 discrete steps (approximately equivalent to an 8mm full-width half-maximums, FWHM) (Hagler et al., 2006), component-based noise correction method to remove motion and physiological artifacts (Behzadi et al., 2007), bandpass filtering (0.008-0.09), and linear detrending.
- Construction of functional connectomes. Functional connectomes were constructed by extracting the BOLD signal from the anatomically defined sub-network, which comprised the 20% of nodes with the largest degree and showing mutual connections (n=9) (Hagmann et al., 2008) in the anatomical data. Fisher’s z-transformed bivariate correlation coefficients were calculated to generate a 9 ⨉ 9 functional connectivity matrix.

## Data analysis

We aimed to investigate whether the structural and functional properties of the brain network can predict the level of coherence in the cultural transmission of auditory codes. First, we used structural connectome analysis to identify an anatomical network whose edge-specific properties relate to TC (**Figure 3**). For that purpose, we used Mrtrix3 software and threshold-free network-based enhanced statistics (TFNBS) (Baggio et al., 2018). Structural connectome matrices were based on the inter-subject normalised connection density (fiber bundle capacity) which is a measure of how much information a white matter pathway can carry. We performed a generalised linear model (GLM) using TC as the main explanatory variable, while accounting for six other factors: age and gender; score in the Gold-MSI subscale for musical training (Müllensiefen et al., 2013), seeding group; head motion during scans (Bastiani et al., 2019), and logarithm (base 10) of the intracranial volume (log-eICV). Edgewise family-wise error corrected (FWE) p-values were obtained with non-parametric permutations (n=5000) (Winkler et al., 2014). Within the resulting network, we selected the 20% of interconnected nodes with the highest degree (N=9). We use these nodes to construct a functional connectivity matrix (9 x 9), where we performed non-parametric spatial pairwise clustering (SPC) (Zalesky et al., 2012). SPC calculates the level of significance in functional connectivity analyses by analysing the connections between brain regions using randomisation or permutation. This method provides cluster-based statistics that are less conservative than those obtained through other methods such as Bonferroni correction, the false discovery rate procedure, and extreme statistics. Using GLM, we tested an association between clusters in the subnetwork and TC scores, while accounting for age and gender, Gold-MSI score, log-eICV, and seeding group.

## Results

Transmission coherence in signalling games. Participants were consistent in reproducing signals and their mappings to meanings throughout Game 2. Transmission Coherence (TC) scores were positive for most participants and significantly different from zero (mean= 0.38, p<0.001). Out of the 51 participants, 47 had a TC-score greater than zero. Only 1 participant had a TC-score equal to zero, indicating no negotiation of code mappings between players; 3 participants had a negative TC-score, indicating a low consistency in using the same code from trial to trial.

Structural connectivity. We identified a subnetwork of 48 cortical nodes connected by 70 edges, which were mostly located in frontal (35%, N=17), temporal (22%, N=11), and insular regions (18%, N=9) and were evenly distributed between the two hemispheres (**Figure 4**) (**Supplementary Tables 1 and 2).** Most edges were fronto-frontal (17%, N=12), followed by frontal-subcortical, frontal-insular (11%, N=8), and fronto-temporal (10%, N=7), and all but two edges were inter-hemispheric. Within this network, the analysis revealed a negative relationship between fiber bundle capacity (FBC) and transmission coherence (TC) scores, meaning that participants with stronger inter-hemispheric connections had lower TC scores, as shown in **Figure 4**.

**Figure 4.**
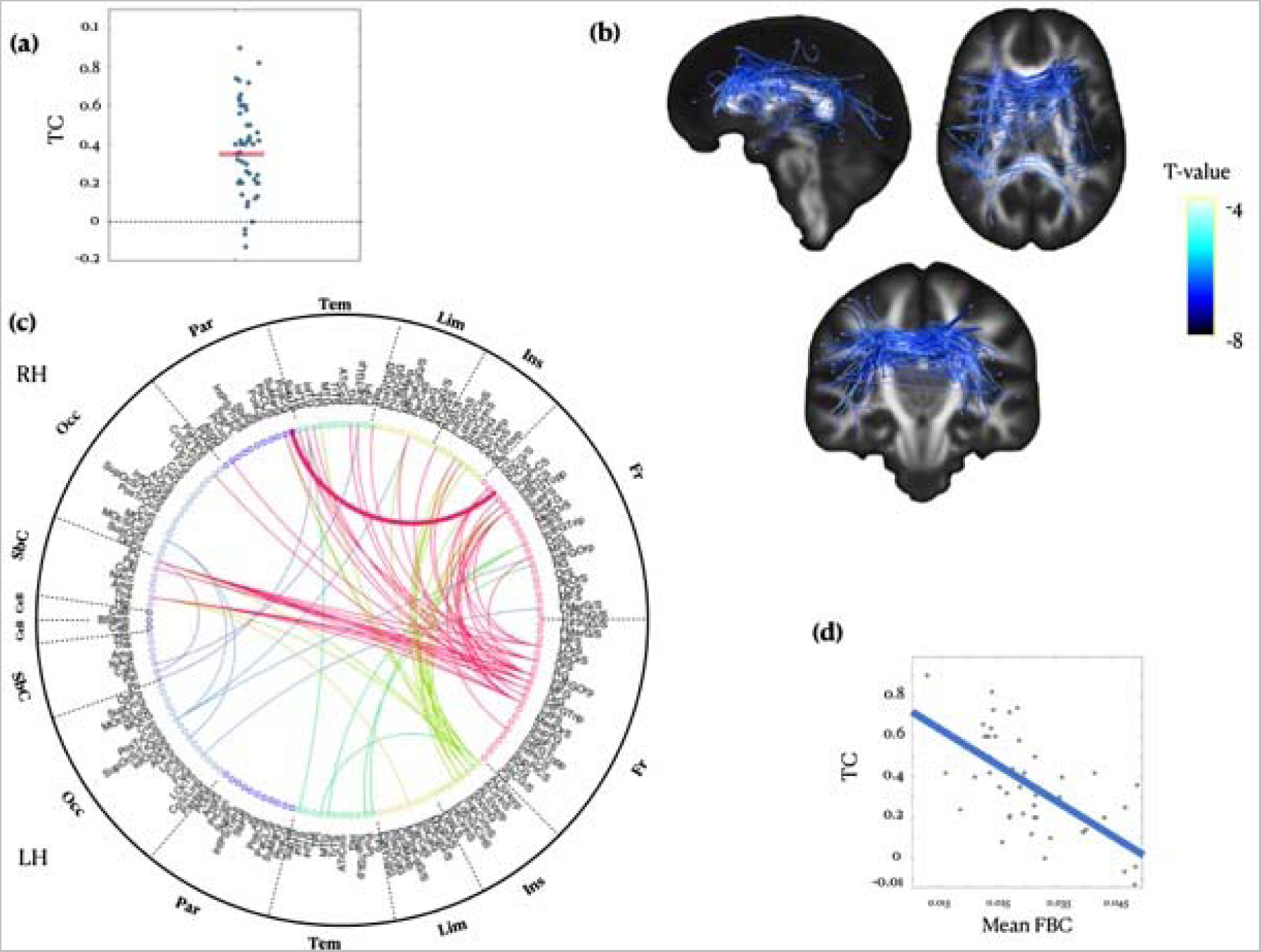
Neuroanatomical network where fiber bundle capacity (FBC) is correlated with transmission coherence in signalling games (Game 2) after controlling for nuisance regressors. (**a**) Scatter plot of TC scores collected in the second game of signalling games. Each point in the scatter plot represents one participant (N = 51). **(b)** Sagittal (top left), axial (top right), and coronal views (bottom) of the structural brain network as revealed by TFNBS (p<0.05, FWE corrected). The colour bar indicates t-values. **(c)** Circular connectogram organized according to brain regions from the Destrieux anatomical atlas (edge-level statistics in Supplementary Table 2). Line colours code for brain lobes, while their size indicates the fiber bundle capacity (FBC). **(d)** Scatter plot showing the negative relationship between mean FBC within the anatomical network and TC scores in signalling games (Game 2). Each point in the scatter plot represents one participant (N = 51).

Functional connectivity. Within the anatomical network, we selected the 20% of nodes with the highest degree (“hubs”) and that were mutually connected (**Supplementary Figure 1**). These nodes were equally distributed across the two hemispheres and belong to the lateral frontal and insular lobes: the orbital part of the left inferior frontal gyrus (InfFGOrp, degree=9), the anterior segment of the circular sulcus of the left insula (ACirInS, degree=7), left inferior frontal sulcus (InfFS, degree=7), left middle frontal gyrus (MFG, degree=6), the opercular part of the right inferior frontal gyrus (InfFGOpp, degree=6), the inferior part of the left precentral sulcus (InfPrCS, degree=5), the left lateral orbital sulcus (LOrS, degree=4), the right subcentral gyrus (central operculum) and sulci (SbCG/S, degree=4), and right short insular gyri (ShoInG, degree=4). Within this network of nine nodes, we performed SPC (**Figure 5**). We found that an increased FC between bilateral inferior frontal areas (right InfPrCS and left InfFS and LOrS) was associated with decreased TC scores (mass=36.08, p_FWE_=0.01).

**Figure 5.**
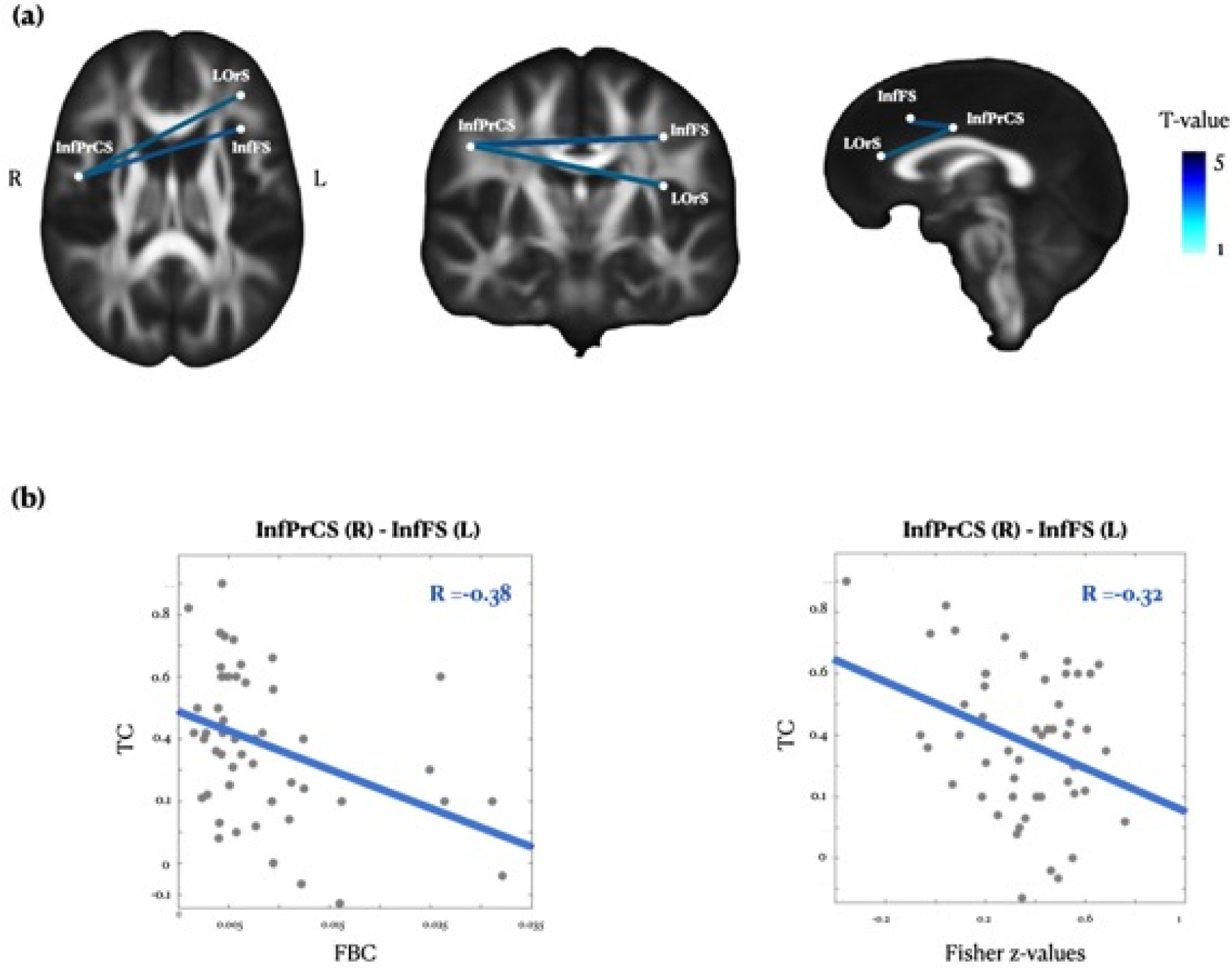
Frontal network where resting-state FC predicts transmission coherence. **(a)** Axial (left), coronal (central), and sagittal views (right) of the resting-state functional network where Fisher-z values were related to TC scores in the signalling games, as revealed by SPC (p<0.05, FWE corrected). **(b)** Scatter plots showing a negative relationship between structural (left) and functional (right) connectivity measures calculated between right inferior precentral sulcus (InfPrCS) and left inferior frontal sulcus (InfFS), and TC scores collected in signalling games (Game 2). Each point in the scatter plot represents one participant (N = 51).

## Discussion

This study combined a laboratory model of cultural transmission, i.e., the signalling games, with neuroimaging methods. In particular, we focused on the sender’s role in the signalling games by measuring the coherence of information transmission from the sender to the receiver. Here, higher coherence was achieved by having a sender transmitting the same signal and maintaining the same signal-meaning mapping throughout the experimental trials. In contrast, lower coherence indicated that the sender introduced variations and errors when composing the signals or mapping them to meanings. We found that senders who exhibited more coherence in the transmission were characterized by weaker inter-hemispheric structural and functional connectivity patterns in the brain. Specifically, higher transmission coherence negatively correlated with the FBC of fronto-frontal, frontal-subcortical, frontal-insular and fronto-temporal edges. From the neuroanatomical network associated to transmission coherence, we extracted a subnetwork of frontal nodes which were characterized by the highest degree and assessed the relationship of coherence with patterns of functional interactions. We found that increased connectivity between bilateral inferior frontal areas (right inferior part of the left precentral sulcus with left inferior frontal sulcus and lateral orbital sulcus) was associated with decreased coherence in the transmission of the information.

Our findings correlate with previous research highlighting relationships between structural and functional features of the brain and complex cognitive tasks. For instance, it has been shown that the integrity of the callosal genu was associated with increased left-lateralization of prefrontal activity during complex cognitive tasks such as encoding of verbal information (Persson et al., 2006; Putnam et al., 2008). Notably, the corpus callosum is the main brain structure responsible for connecting left and right hemispheres. Thus, reduced size of corpus callosum suggests a lower degree of connectivity and communication between regions of the two hemispheres and hence a higher of lateralization in their neural processes. This could plausibly explain that in our study, coherence across trials in reproducing melodies and their mappings to meaning was correlated negatively to functional and structural connectivity between hemispheres, indicating that reduced inter-hemispheric connectivity may be beneficial for coherent transmission of auditory information. Another study focusing on the corpus callosum discovered that the size of the rostral corpus callosum was associated with the brain processes underlying memory abilities (Kompus et al., 2011). The authors found that a larger genu predicted more pronounced voxel-based asymmetry during memory processes such as encoding and retrieval of face-name mappings. Notably, this suggested that, even when inter-hemispheric connections are enhanced, complex cognitive tasks may still rely on a higher degree of lateralization in the brain. Since some amount of memory abilities is required in the signalling games used in our study, the results reported by Kompus et al. (2011) are in line with our findings. In fact, Kompus et al. (2011) suggested that increased connections in the rostral corpus callosum promoted communication between homologous frontal regions which may not be favourable during a complex memory task relying on hemispheric lateralization. Similarly, we observed that increased connectivity between bilateral inferior frontal areas (right inferior part of the left precentral sulcus with left inferior frontal sulcus and lateral orbital sulcus) was associated with decreased coherence in the transmission of the information.

Along this line, additional previous works highlighted that lateralization of brain processing can be beneficial for complex cognitive functions. In an fMRI study, Golby et al. (2001) studied the brain lateralization during encoding of information. The authors found that verbal encoding produced left-lateralized activation of the inferior prefrontal cortex and the medial temporal lobe (MTL), while pattern encoding activated the right inferior prefrontal cortex and right MTL. Their results showed that the two hemispheres were specialized in encoding different types of information suggesting that a more specialized encoding process may benefit from more hemispheric segregation. Similarly, our study revealed decreased inter-hemispheric connectivity in individuals who exhibited higher coherence in the transmission of information; a complex process which largely implies encoding of information. Since a decreased inter-hemispheric connectivity suggests a higher tendency towards hemispheric lateralization, our results are very much in line with the findings reported by Golby et al. (2001).

In the context of auditory processing of information, several studies showed that the auditory cortex (AC) exhibited functional asymmetry between the left and right hemispheres (Toga & Thompson, 2003). Verbal sound processing such as of speech predominantly occurred in the left hemisphere, while nonverbal sound processing, such as noise, was dominant in the right hemisphere (Tervaniemi & Hugdahl, 2003). The extent of hemispheric lateralization was affected by the surrounding environment (Tervaniemi & Hugdahl, 2003). For instance, in a quiet environment, the left AC was more involved in speech processing, but in a noisy environment, the recruitment of right AC was more pronounced (Santosa et al., 2014; Tervaniemi & Hugdahl, 2003), suggesting that lateralization played a critical role in processing complex information. Another key theory in the functional lateralization of auditory and speech information processing is the ‘asymmetric sampling in time’ (AST) hypothesis proposed by Poeppel, which states that auditory processing occurs by means of anatomic symmetry and functional asymmetry (Giraud & Poeppel, 2012; Giroud et al., 2020; Poeppel, 2003). This hypothesis is built on two key observations: 1) speech signals contain multiple time scales which are relevant to auditory cognition (e.g., time scales related to processing transitions of formants compared to scales associated with syllabicity and intonation); 2) the perception of speech is processed by both left and right auditory cortices with inter-hemispheric similarity and differences. Building on those observations, AST states that the input signal of speech has a symmetric inter-hemispheric neural representation at an early representational level. After that, the signal is processed in an asymmetric manner with regards to the time domain. Here, left auditory regions extract information from short (∼20–40 ms) time-windows. Conversely, the right hemisphere of the brain processes information from later and longer (∼150–250 ms) time-windows. The AST hypothesis reinforces the idea that complex extraction and elaboration of auditory information happens by lateralizing different processes. Coherently with these investigations on auditory perception, in our study we found that coherent transmission of information using five-tone auditory signals was enhanced in participants characterized by a reduced inter-hemispheric connectivity.

Finally, Caeyenberghs and Leemans (2014) investigated the brain structural connectivity of the two hemispheres of a large sample of over 300 participants. They found that the left hemisphere exhibited greater efficiency compared to the right one, whereas the right hemisphere showed higher levels of betweenness centrality and small-worldness, measures which indicate the efficiency of a network. Specifically, brain regions related to language and motor actions displayed increased nodal efficiency in left-hemispheric networks, whereas regions involved in memory and visuospatial attention exhibited increased nodal efficiency in the right hemisphere. These findings suggest that complex cognitive tasks may require a certain degree of lateralization and that, if there is more inter-hemispheric connectivity, the efficiency of information processing in the brain may be reduced. In this light, it is not surprising that our results pointed to a positive correlation between decreased inter-hemispheric connectivity and increased coherence in information transmission.

Taken together, these findings support the hypothesis that hemispheric lateralization may be beneficial for performing a variety of complex cognitive tasks. Interestingly, in our study we found that most of the connections which correlated with decreased coherence of information transmission were inter-hemispheric and involved frontal regions. Thus, if coherence in transmission depends on lateralized engagement of frontal regions rather than on an equal division of inter-hemispheric labor, larger inter-section size or greater density of fiber bundles would have a detrimental effect on transmission of the information. Hence, our results may indicate that to preserve coherence in the information transmission, a certain degree of brain lateralization may be required.

Decreased coherence in information transmission leads to worse replication of the original message. However, this should not only be considered as a negative event. In fact, to allow cultural evolution and development, a particular level of innovation in the interactions between individuals is essential (Cavalli-Sforza & Feldman, 1981; Cavalli-Sforza et al., 1982; Mesoudi & Whiten, 2008). In this light, our results open new perspectives for understanding the neural underpinnings of cultural transmission, creativity, and, in a wider view, cultural divergence and innovation. More specifically, it may be argued that increased inter-hemispheric connectivity is crucial for the process of divergence of information. This is in line with previous studies which suggested that creativity may rely on long-range neural connections which enhance communication between distributed networks (Heilman, 2016; Travis, 2021) and integrate sensory-motor information (Kenett et al., 2018). For instance, in an fMRI study, Villarreal et al. (2013) measured the brain activity of high and low creative participants while they improvised on different rhythmic fragments. Individuals with greater creative abilities derived from increased fluidity and flexibility of their performance displayed enhanced activity in the prefrontal brain regions of both hemispheres as well as in the right insula. Conversely, those with poorer abilities made only slight changes to the original rhythms and exhibited activity only in the left unimodal areas of the brain. In line with our results, this study showed that reproduction of the same acoustic material was associated with hemispheric segregation, while divergence and creativity required the recruitment of both brain hemispheres.

Although our findings provide novel insights into the neural mechanisms associated to coherent transmission of information, the current study has a few limitations that should be discussed. First, our investigation did not involve any functional task in the scanner and thus could not study the lateralization of the brain while participants performed the signalling games. The support of hemispheric lateralization was instead inferred indirectly by measuring the strength of resting-state functional and structural inter-hemispheric connections. For this reason, even if our results provide reliable evidence on the relationship between brain connectivity and transmission of information, future works are required to study the brain activity specifically associated to signalling games. Second, in our study, most measures of information transmission were positive, indicating that participants (senders) transmitted the code coherently throughout the game. Thus, we cannot draw any conclusion on the relationship between the information transmission primarily determined by the receivers and their inter-hemispheric connectivity. Conversely, we can state that senders who transmitted coherent information throughout the experiment exhibited less inter-hemispheric connections. Hence, more research is needed to fully determine the structural and functional neural underpinnings of information transmission determined by the receivers in the signalling games.

In conclusion, our study focused on coherent transmission of information, highlighting some of the neural mechanisms that may be responsible for coherence and divergence in cultural evolution. These findings integrated and expanded on our previous work on the neural basis of cultural transmission which highlighted that neurophysiological components and brain network measures were able to predict performances in signalling games (M. Lumaca et al., 2021; Lumaca et al., 2023; Massimo Lumaca et al., 2021; Lumaca, Haumann, et al., 2018; Lumaca, Kleber, et al., 2019; Lumaca, Ravignani, et al., 2018; Lumaca, Trusbak Haumann, et al., 2019; Lumaca et al., 2022). Taken together, our studies provided an extensive picture of the brain structures arguably required for cultural transmission; however, they did not reveal the fast-scale spatiotemporal dynamics of the brain regions directly involved in the transmission, reception, and interpretation of the information. Thus, future studies are called to investigate the specific neural dynamics underlying coherent transmission of information as measured in laboratory models such as signalling games using a combination of neurophysiology (e.g. magnetoencephalography and electroencephalography) and neuroimaging (e.g. fMRI).

## Authors contribution

ML conceived the hypotheses and designed the study. ML, PV and LB recruited the resources for the experiment and writing up the manuscript. ML, AKV and CI collected the data. ML and AKV performed pre-processing and statistical analysis. ML, LB and PV provided essential contribution to interpret and frame the results within the neuroscientific literature. LB and ML wrote the first draft of the manuscript. ML prepared the figures. All the authors contributed to and approved the final version of the manuscript.

