## Supplementary Material for "Decreased inter-hemispheric connectivity predicts a coherent retrieval of auditory symbolic material in a laboratory model of cultural transmission"

**1. Instructions**

**1.1. Session 1 - MRI**

The MRI session took place on a CE-approved 3T Siemens MR-scanner at Neuroradiologisk forskningsenhed, AUH, Nørrebrogade 44, 8000 Århus C. Before the start of the session, the investigator ensured that the participant presented no contra-indications to an MR examination. This was performed by going through a list of questions with the participant, detailed in an MRI screening form. All participants were informed that they could discontinue participating in the experiment at any time. Once the written information consent was signed and the MRI screening form completed, the participant was prepared for the experiment. The experimenter asked the participant to remove any wearable metal items and explained the instructions of the study. The participant was then installed on the bed of the MR scanner and equipped with the necessary stimulation equipment (earplugs and stereo headphones to reduce the scanner noise) and recording equipment (ECG and respiration). An emergency call button was also given to the participant to alert the experimenter during scanning, if necessary. Once the participant was equipped, the scanner bed was entered into the magnet bore and the experiment started. The MRI session was divided into 3 main scans. The first scan was resting state (rsFMRI; 9 min). We asked the participant to relax inside the MRI scanner without engaging in any specific task. During the second scan (fMRI oddball paradigm; 25 min), auditory stimulation was delivered to the participant via headphones in 3 separate blocks (for details see Lumaca et al., 2019). The third scan was structural (T1 and DTI; 20 min). During the structural scan the participants were asked to stay steady and to watch a documentary movie projected on an MRI-compatible display. Throughout the examination, visual contact (through the glass panel of the examination room) and auditory contact (through a microphone in the scanner bore) were kept with the participant to ensure normal unfolding of the experimental session. In the meanwhile, brain scans were acquired. Once the scan acquisition was finished, the participant was taken out of the scanner and asked for feedback. The time of scan acquisition did not exceed 60 minutes (plus 30 min of information, preparation and debriefing).

**1.2. Session 2 - Signaling Games**

**Participant**

Before the signaling games (SGs) session, each participant was informed that they would take part in two successive games, each played in a different role. In Game 1, they would be the receiver. They were asked to try to learn which facial expressions (emotions) corresponded to the 5-equitone sequences sent by the other player (sender). There was no reward for speed, either in a trial or in the game. Successful coordination was the only reward for participants. In Game 2, the participant played as sender. They were instructed to transmit the tone system acquired in Game 1, as they remembered it, to a new receiver.

**Confederate**

**Game 1.** The confederate was trained on a particular mapping of emotions to keys on the keyboard (see Procedure, main article): in Game 1, they played with no sound feedback (volume set at 0), to prevent him from learning the ‘seed’ system and be biased in Game 2. The confederate was instructed to use consistently that mapping throughout Game 1, regardless of what the participant (receiver) would do.

**Game 2.** The confederate played according to the same written instructions given to the participant (receiver) in Game 1, namely, they were instructed to try to learn the temporal patterns as produced by the participant (sender).

**2. Supplementary Figures**

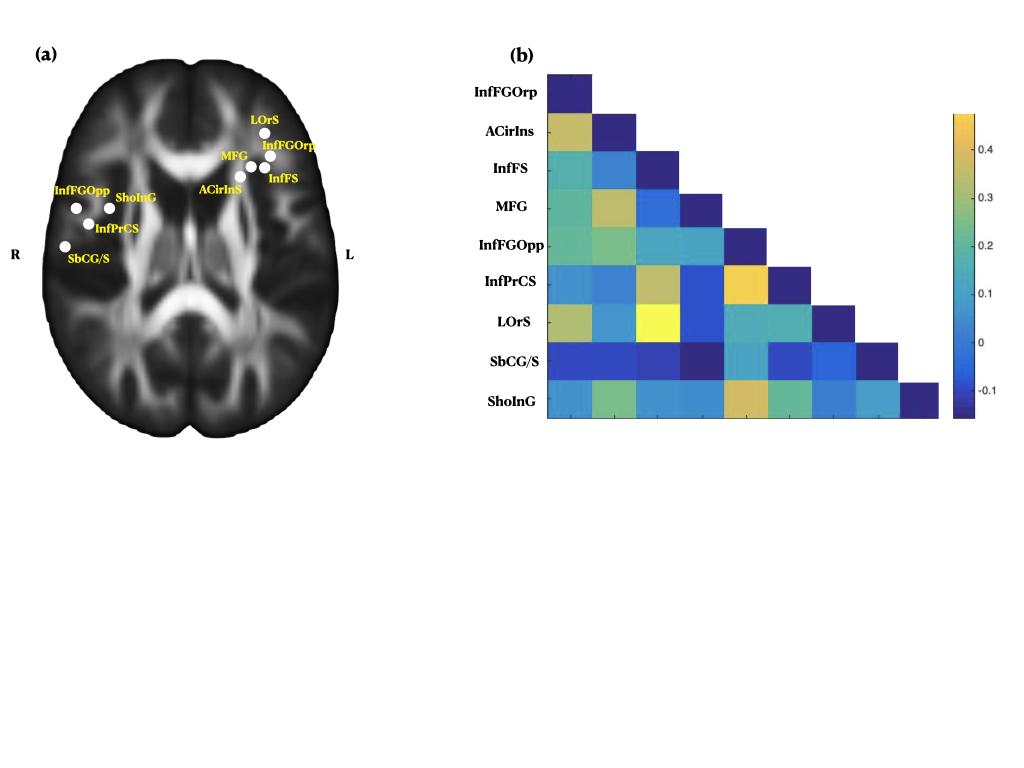

**Supplementary Figure 1.** Subnetwork of regions-of-interest (ROIs) where rs-FC analysis was performed. (a) These brain regions were selected from the structural connectome-based analysis: they show the highest degree and share mutual connections. (b) Functional connectivity matrix of the anatomical subnetwork. Bars represent the Fisher z-transformed correlation coefficients.

**3. Supplementary Tables**

**Supplementary Table 1.** Abbreviations and descriptions of the brain structures (48 cortical and 5 subcortical) showing significant edges within connectome group-wise analyses. Prefixes: l=left; r=right.

| **Abbreviation** | **Description** | **Lobe** |
| --- | --- | --- |
| *Neocortical structures* | | |
| lPosTrCoS | Posterior transverse collateral sulcus | Occipital |
| lAOcS | Anterior occipital sulcus and preoccipital notch (temporo-occipital incisure) |  |
| lFuG | Lateral occipito-temporal gyrus (fusiform gyrus, O4–T4) |  |
| rMOcS/LuS | Middle occipital sulcus and lunatus sulcus |  |
| rCoS/LinS | Medial occipito-temporal sulcus (collateral sulcus) and lingual sulcus |  |
| rAngG | Angular gyrus | Parietal |
| rSuMarG | Supramarginal gyrus |  |
| rPosCG | Postcentral gyrus |  |
| lSupTS | Superior temporal sulcus | Temporal |
| rSupTS |  |  |
| lTPl | Temporal plane of the superior temporal gyrus |  |
| rTPl |  |  |
| lMTG | Middle temporal gyrus (T2) |  |
| lPoPl | Polar plane of the superior temporal gyrus |  |
| lTPo | Temporal pole |  |
| rTPo |  |  |
| rInfTS | Inferior temporal sulcus |  |
| rInfTG | Inferior temporal gyrus (T3) |  |
| rHG | Heschl's gyrus (anterior transverse temporal gyrus) |  |
| rSupTGLp | Lateral aspect of the superior temporal gyrus |  |
| lPerCaS | Pericallosal sulcus (S of corpus callosum) | Limbic |
| rPerCaS |  |  |
| lSupCirIns | Superior segment of the circular sulcus of the insula | Insula |
| rSupCirIns |  |  |
| lShoInG | Short insular gyri |  |
| rShoInG |  |  |
| lALSVerp | Vertical ramus of the anterior segment of the lateral sulcus (or fissure) |  |
| rALSVerp |  |  |
| lACIrIns | Anterior segment of the circular sulcus of the insula |  |
| lALSHorp | Horizontal ramus of the anterior segment of the lateral sulcus (or fissure) |  |
| rPosLS | Posterior ramus (or segment) of the lateral sulcus (or fissure) |  |
| lInfFGopp | Opercular part of the inferior frontal gyrus | Frontal |
| rInfFGTopp |  |  |
| lInfFS | Inferior frontal sulcus |  |
| lInfFGTrip | Triangular part of the inferior frontal gyrus |  |
| rInfFGTrip |  |  |
| lOrG | Orbital gyri |  |
| lMFG | Middle frontal gyrus (F2) |  |
| rMFG |  |  |
| lInfFGOrp | Orbital part of the inferior frontal gyrus |  |
| rInfFGOrp |  |  |
| lOrS | Lateral orbital sulcus |  |
| lLOrS | Lateral orbital sulcus |  |
| rFMarG/S | Fronto-marginal gyrus (of Wernicke) and sulcus |  |
| rInfPrCS | Inferior part of the precentral sulcus |  |
| rSupPrCS | Superior part of the precentral sulcus |  |
| rPrCG | Precentral gyrus |  |
| SbCG/S | Subcentral gyrus (central operculum) and sulci |  |
| *Subcortical structures* | | |
| lCaN | Caudate nucleus |  |
| lPal | Pallidum |  |
| rPal |  |  |
| rTha | Thalamus |  |
| rPu | Putamen |  |

**Supplementary Table 2.** Edgewise statistics for the structural brain network identified with TFNBS (p_FWE_<0.05). Lobe-to-lobe connections (N=70) are arranged by number in descending order. Prefixes: l=left; r=right.

| **Abbreviation(s)** | | **Effect size^a^** | **T-statistic (p_FWE_)** |
| --- | --- | --- | --- |
| *Fronto-frontal (inter-hemispheric)* | | | |
| InfFGOrp | InfFGTopp | 2.62 | 4.41 (0.008) |
|  | MFG | 2.21 | 3.52 (0.03) |
| InfFGTopp | InfPrCS | 2.20 | 3.51 (0.03) |
|  | InfFGTopp | 2.18 | 3.47 (0.03) |
|  | InfPrCS | 2.38 | 3.79 (0.03) |
| InfFS | SbCG/S | 2.83 | 4.50 (0.009) |
|  | InfFGTopp | 2.16 | 3.45 (0.03) |
| InfFS | InfPrCS | 2.14 | 3.42 (0.03) |
| LOrS | sbCG/S | 2.34 | 3.72 (0.02) |
|  | InfFGTopp | 2.52 | 4.01 (0.01) |
| MFG | SbCG/S | 2.19 | 3.48 (0.03) |
| OrS | SupPrCS | 2.33 | 3.70 (0.02) |
| *Frontal-subcortical (interhemispheric)* | | | |
| InfFGOrp | Tha | 2.59 | 4.12 (0.008) |
|  | Pu | 2.72 | 4.33 (0.007) |
|  | Pal | 2.52 | 4.02 (0.01) |
| InfFS | Tha | 2.36 | 3.75 (0.02) |
|  | Pu | 2.36 | 3.76 (0.02) |
| LOrS | Pu | 2.37 | 3.78 (0.02) |
| MFG | Tha | 2.46 | 3.91 (0.01) |
|  | Pu | 2.25 | 3.58 (0.03) |
|  | Pal | 2.28 | 3.63 (0.02) |
| OrG | Tha | 2.15 | 3.42 (0.03) |
|  | Pu | 2.53 | 4.03 (0.01) |
| OrS | Pu | 2.65 | 4.22 (0.007) |
| *Frontal-temporal (inter-hemispheric)* | | | |
| OrG | HG | 2.21 | 3.51 (0.03) |
|  | SupTS | 2.21 | 3.52 (0.03) |
| OrS | TPo | 2.83 | 4.51 (0.005) |
|  | infTS | 2.26 | 3.60 (0.02) |
| MFG | SupTGLp | 2.85 | 4.53 (0.007) |
| MTG | InfFGTrip | 2.30 | 3.65 (0.02) |
| InfFGOrp | infTG | 2.17 | 3.45 (0.03) |
| InfFGTrip | TPo | 2.59 | 4.13 (0.02) |
| PoPI | InfFGOrp | 2.93 | 4.67 (0.02) |
| *Frontal-insular (inter-hemispheric)* | | | |
| InfFGOrp | ShoInG | 2.54 | 4.04 (0.01) |
|  | SupCirlnS | 2.61 | 4.15 (0.008) |
| InfFGTopp | ACirlnS | 2.58 | 4.10 (0.008) |
| InfFGTopp | ALSVerp | 2.18 | 3.48 (0.03) |
| InfPrCS | ALSVerp | 2.31 | 3.67 (0.03) |
| InfFS | PosLS | 2.18 | 3.47 (0.03) |
| SbCG/S | ShoInG | 2.18 | 3.47 (0.03) |
| InfFGTrip | ShoInG | 2.22 | 3.53 (0.03) |
| *Frontal-occipital (inter-hemispheric)* | | | |
| InfFS | CoS/LinS | 2.48 | 3.95 (0.01) |
| MFG | PosTrCos | 2.30 | 3.66 (0.02) |
| FMarG/S | FuG | 2.47 | 3.93 (0.03) |
| *Insular-insular (inter-hemispheric)* | | | |
| ACirlnS | ShoInG | 2.48 | 3.94 (0.01) |
| ACirlnS | ALSVerp | 2.47 | 3.94 (0.01) |
|  | SupCirlnS | 2.20 | 3.50 (0.03) |
| ALSHorp | ShoInG | 2.37 | 3.78 (0.02) |
| *Insular-temporal (inter-hemispheric)* | | | |
| ALSHorp | TPl | 2.17 | 3.46 (0.03) |
| *(intra-hemispheric)* | | | |
| ALSVerp | TPl | 2.19 | 3.49 (0.03) |
| *Frontal-parietal (inter-hemispheric)* | | | |
| MFG | AngG | 2.21 | 3.53 (0.03) |
| OrG | SuMarG | 2.19 | 3.50 (0.03) |
| *(intra-hemispheric)* | | | |
| PrCG | PosCG | 2.19 | 3.49 (0.04) |
| *Insular-Subcortical (inter-hemispheric)* | | | |
| ACirlnS | Tha | 2.37 | 3.77 (0.02) |
|  | Pu | 2.68 | 4.27 (0.007) |
| *Insular-Parietal (inter-hemispheric)* | | | |
| ACirlnS | SuMarG | 2.26 | 3.60 (0.02) |
| ALSHorp | SuMarG | 2.22 | 3.54 (0.03) |
| *Frontal-temporal (inter-hemispheric)* | | | |
| LOrS | SupTS | 2.52 | 4.01 (0.01) |
| *Temporo-temporal (inter-hemispheric)* | | | |
| SupTS | InfTS | 2.19 | 3.48 (0.03) |
| *Subcortical-temporal (inter-hemispheric)* | | | |
| CaN | SupTGLp | 2.44 | 3.88 (0.01) |
| Pu | inFTS | 2.16 | 3.44 (0.03) |
| *Occipito-occipal (inter-hemispheric)* | | | |
| AOcS | MOcS/LuS | 2.14 | 3.41 (0.03) |
| PosTrCoS | MOcS/LuS | 2.30 | 3.67 (0.02) |
| *Subcortical-subcortical (inter-hemispheric)* | | | |
| Pal | Pu | 2.25 | 3.58 (0.03) |
| *Limbic-insular* | | | |
| PerCaS | ALSHorp | 2.21 | 3.52 (0.03) |
| *Limbic-frontal* | | | |
| PerCaS | InfFGOrp | 2.14 | 3.41 (0.03) |
| *Limbic-occipital* | | | |
| PerCaS | PosTrCoS | 2.32 | 3.69 (0.02) |
| *Limbic-subcortical* | | | |
| PerCaS | Tha | 2.23 | 3.56 (0.03) |

^a^Effect size is Cohen’s D
